# Conducting social network analysis with animal telemetry data: applications and methods using spatsoc

**DOI:** 10.1101/447284

**Authors:** Alec L. Robitaille, Quinn M.R. Webber, Eric Vander Wal

## Abstract

1. We present `spatsoc`: an R package for conducting social network analysis with animal telemetry data.
2. Animal social network analysis is a method for measuring relationships between individuals to describe social structure. Using animal telemetry data for social network analysis requires functions to generate proximity-based social networks that have flexible temporal and spatial grouping. Data can be complex and relocation frequency can vary so the ability to provide specific temporal and spatial thresholds based on the characteristics of the species and system is required.
3. `spatsoc` fills a gap in R packages by providing flexible functions, explicitly for animal telemetry data, to generate gambit-of-the-group data, perform data-stream randomization and generate group by individual matrices.
4. The implications of spatsoc are that current users of large animal telemetry or otherwise georeferenced data for movement or spatial analyses will have access to efficient and intuitive functions to generate social networks.

## Introduction

Animal social network analysis is a method for measuring the relationships between individuals to describe social structure (Wey et al. 2008; Croft, James, and Krause 2008). Association networks are built from a set of observed elements of social community structure and are useful to understand a variety of ecological and behavioural processes, including disease transmission, interactions between individuals and community structure (Pinter-Wollman et al. 2014). Among the most common types of social network data collection is gambit-of-the-group, where individuals observed in the same group are assumed to be associating or interacting (Franks, Ruxton, and James 2010). Similar to gambit-of-the-group, proximity based social networks (PBSNs) are association networks based on close proximity between individuals (Spiegel et al. 2016). PBSNs rely on spatial location datasets that are typically acquired by georeferenced biologging methods such as radio-frequency identification tags, radiotelemetry, and Global Positioning System (GPS) devices (hereafter, animal telemetry).

Biologging using GPS devices allow simultaneous spatiotemporal sampling of multiple individuals in a group or population, thus generating large datasets which may otherwise be challenging to collect. The advent of biologging technology allows researchers to study individuals of species that range across large areas, migrate long distances, or spend time in inaccessible areas (Cagnacci et al. 2010; Cooke et al. 2013; Hebblewhite and Haydon 2010). Moreover, the recent increase in the number of studies using GPS telemetry to study movement ecology (Kays et al. 2015; Tucker et al. 2018) indicates the potential for a large number of existing datasets that may be retro-actively analyzed to test a priori hypotheses about animal social structure. As animal telemetry data have become more accessible and finer scaled, a number of techniques and methods have been developed to quantify various aspects of animal social structure. These include dynamic interaction networks (Long et al. 2014), PBSNs (Spiegel et al. 2017) and the development of traditional randomization techniques to assess non-random structure of PBSNs constructed using animal telemetry data (Spiegel et al. 2016). Despite the recent increase in the number of studies using animal telemetry data and GPS relocation data (Webber and Vander Wal 2018), there is no comprehensive R package that generates PBSNs using animal telemetry data.

Here, we present, `spatsoc` a package developed for the R programming language (R Core Team 2018) to (i) convert animal telemetry data into gambit-of-the-group format to build PBSNs, (ii) implement data-stream social network randomization methods of animal telemetry data (Farine and Whitehead 2015; Spiegel et al. 2016), and (iii) provide flexible spatial and temporal grouping of individuals from large datasets. Animal telemetry data can be complex both temporally (e.g., data can be partitioned into monthly, seasonal or yearly segments) and spatially (e.g., subgroups, communities or populations). Functions in spatsoc were developed taking these complexities into account and provide users with flexibility to select relevant parameters based on the biology of their study species and systems and test the sensitivity of results across spatial and temporal scales.

## Functions

The `spatsoc` package provides functions for using animal telemetry data to generate PBSNs. Relocations are converted to gambit-of-the-group using grouping functions which can be used to build PBSNs. Raw data streams can be randomized where animal telemetry data is swapped between individuals at hourly or daily scales (Farine and Whitehead 2015), or within individuals using a daily trajectory method (Spiegel et al. 2016).

## Grouping

Gambit-of-the-group data is generated from animal telemetry data where individuals are grouped based on temporal and spatial overlap (Figure 1). The spatsoc package provides one temporal grouping function:

1. `group_times` groups animal telemetry relocations into time groups. The function accepts date time formatted data and a temporal threshold argument. The temporal threshold argument allows users to specify a time window within which relocations are grouped, for example 5 minutes, 2 hours or 10 days. We recommend this temporal threshold is based on the nuances of the animal telemetry data, study species and system.

**Figure 1:**
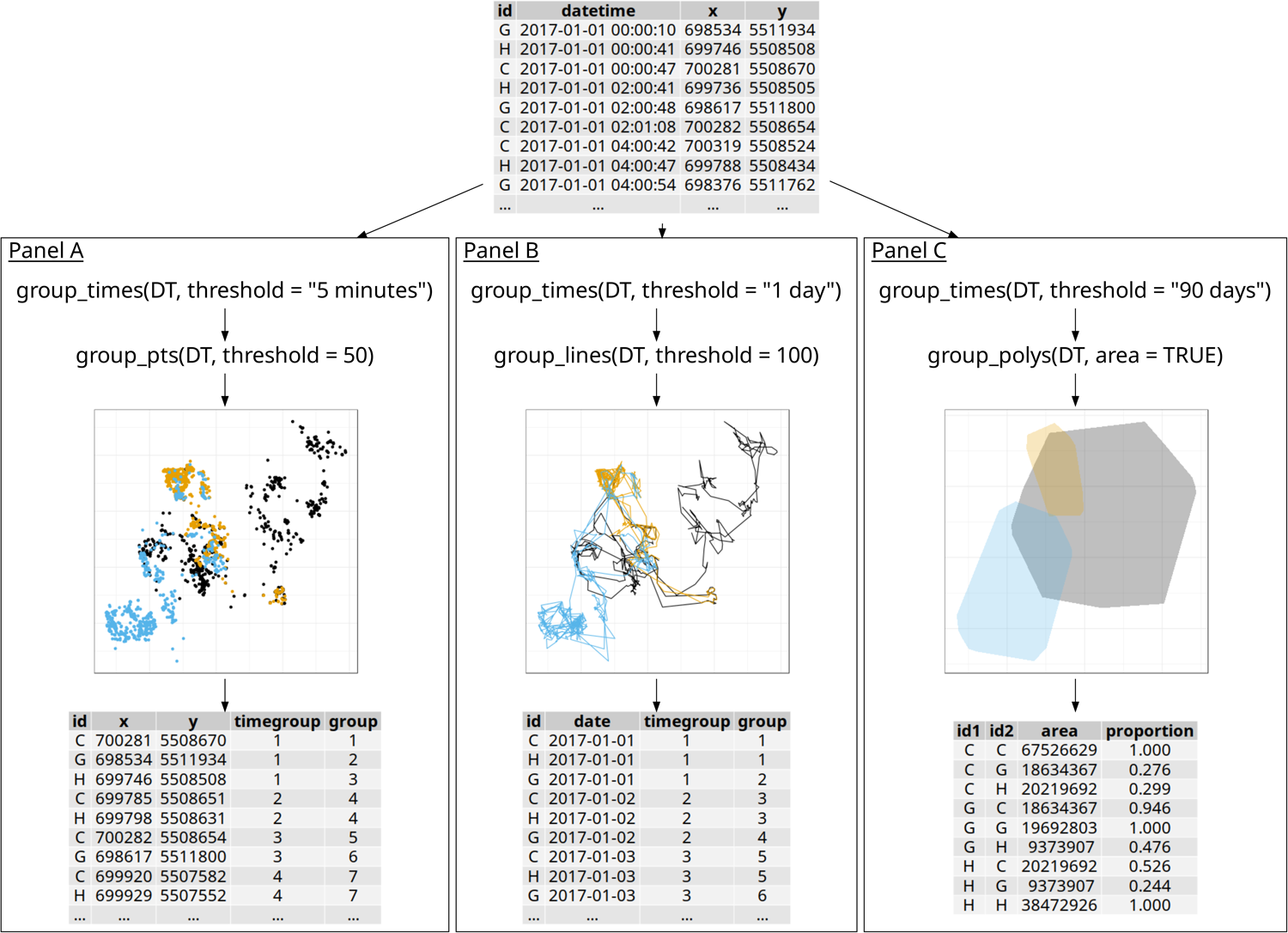
Broad overview flowchart of recommended usage of grouping functions. A temporal grouping function (`group_times`) is followed by one of three spatial grouping functions. Panel A) `group_pts` compares the Euclidean distance between animal telemetry relocations and groups points based on a threshold distance with an output containing temporal and spatial groups for each individual at each relocation; Panel B) `group_lines` groups overlapping movement trajectories generated from GPS relocations with an output containing temporal and spatial groups for each individual at each relocation; or Panel C) `grop_polys` generates and groups overlapping home ranges of individuals and optionally returns a measure of proportional area overlap with an output containing the area of overlap (*m*^2^) and proportion of overlap among individuals. Note, coordinates for each panel are projected in Universal Transverse Mercator and the unit for distance thresholds is in meters.

The `spatsoc` package provides three spatial grouping functions:

1. `group_pts` compares the Euclidean distance between animal telemetry relocations (Figure 2 - Panel A). Relocations for all individuals within each time group will be grouped based on spatial proximity. Spatial proximity is defined by the user-specified distance threshold.
2. `group_lines` groups overlapping movement trajectories generated from animal telemetry data. Movement trajectories for each individual within each time group, e.g. 8 hours, 1 day or 20 days, are generated and grouped based on spatial overlap, optionally within the user-specified distance threshold buffers around each trajectory (Figure 2 - Panel B).
3. `group_polys` generates and groups overlapping home ranges using kernel utilization distributions or minimum convex polygons generated in `adehabitatHR` of individuals and optionally returns a measure of proportional area overlap.

**Figure 2:**
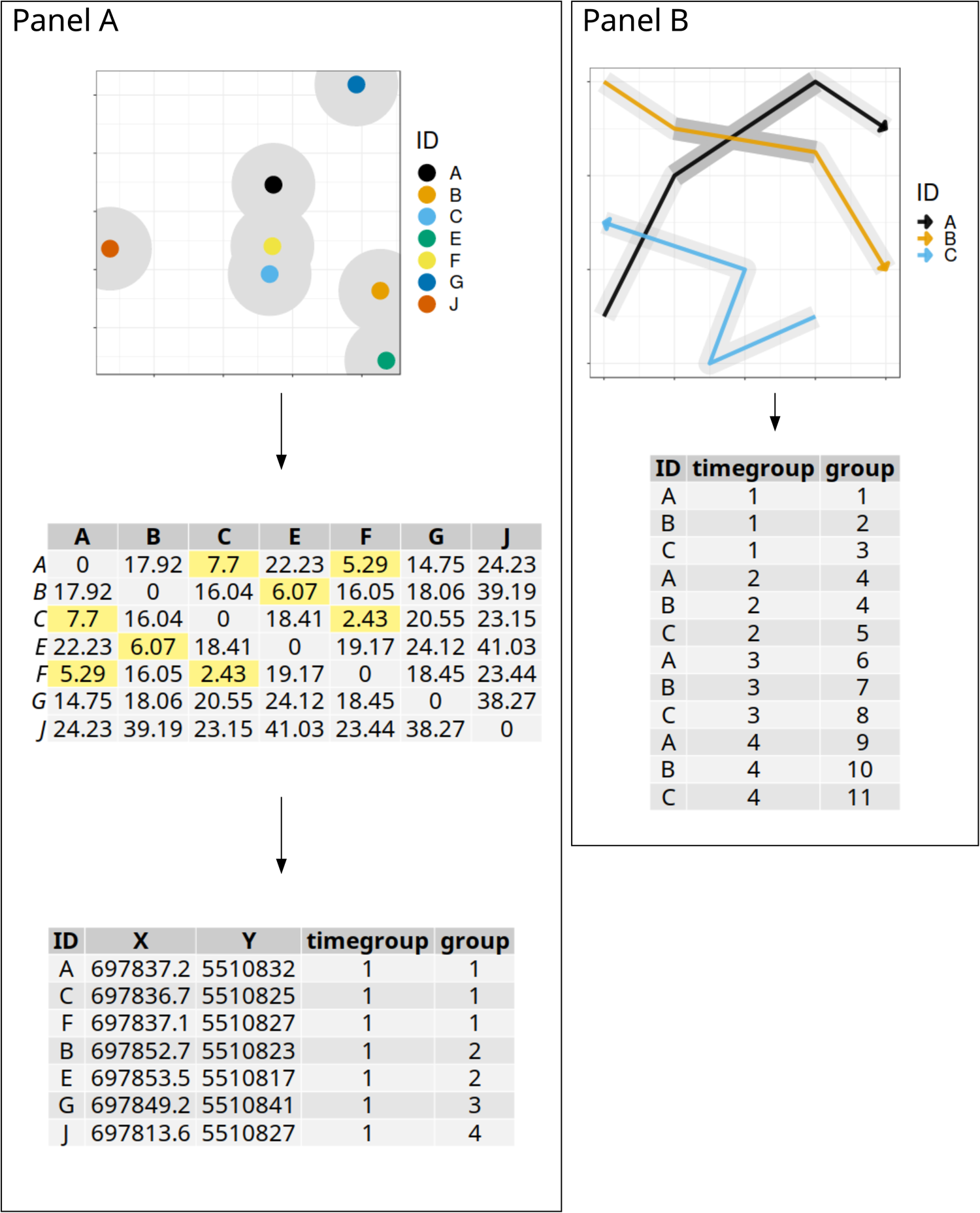
Detailed overview flowchart of `group_pts` and `group_lines`. Panel A) `group_pts` details showing a single spatial relocation for each of seven individuals at the same time, each indicated by a different colour. Based on the distance among spatial relocations, a distance matrix is generated. Distances are compared to the user-provided spatial distance threshold and individuals within this distance are grouped together. The distance threshold in this example is 15 m and distances less than this threshold are highlighted in yellow. Note, *individual J* was not within 15 meters of any other individuals, but is still assigned a group number as a solitary individual. Panel B) `group_lines` details showing buffered lines for three individuals and resulting spatial groups. Within each time group generated by `group_times`, line segments are generated and buffered by the distance threshold and individuals with overlapping buffers, indicated by the dark grey shaded areas, are grouped together. Note, coordinates for each panel are projected in Universal Transverse Mercator and the unit for distance thresholds is in meters.

For all three spatial grouping functions, individuals that are not within the user-specified distance threshold or that do not overlap with any other individuals are assigned to a group on their own. Functions provided by `spatsoc` emphasize flexibility to allow users the ability to modify functions to better suit their study systems. The temporal threshold argument of `group_times` accepts units of minutes, hours and days to consider spatial grouping at different temporal scales. For example, grouping trajectories with `group_lines` is compatible with daily or weekly time group thresholds while point grouping with `group_pts` is compatible with minute or hourly time group thresholds.

## Randomizations

Randomization procedures in social network analysis are important to test assumptions of spatial and temporal non-independence of social association data (Farine and Whitehead 2015). Data-stream randomization is the recommended randomization technique for social network users (Farine and Whitehead 2015) and involves swapping individuals and group observations within or between temporal groups and individuals. Animal telemetry data has inherent temporal structure and is well suited to randomization methods. The `spatsoc` package provides three data-stream randomization methods:

1. Step - randomizes identities of animal telemetry relocations between individuals within each time step.
2. Daily - randomizes daily animal telemetry relocations between individuals, preserving the order of time steps.
3. Trajectory - randomizes daily trajectories generated from animal telemetry relocations within individuals (Spiegel et al. 2016).

The `randomizations` functions return the input data with random fields appended, ready to use by the grouping functions or to build social networks. Step and daily methods return a “randomID” field that can be used in place of the ID field and the trajectory method returns a “randomDatetime” that can be used in place of the datetime field. The function `randomizations` in `spatsoc` allow users to split randomizations between spatial or temporal subgroups to ensure that relocations are only swapped between or within relevant individuals.

## Using spatsoc in social network analysis

`spatsoc` is integrated with social network analysis in R in three main steps: 1) generate gambit-of-the-group data by temporal and spatial grouping, 2) generate group by individual matrices and 3) PBSN data-stream randomization. Users should first determine relevant temporal and spatial grouping thresholds based on details from their study species and systems. These thresholds depend on the fix rate of animal telemetry devices, movement rates of study species, and other biological details of each species and system as well as the questions or hypotheses of interest. In addition, thresholds selected for temporal and spatial grouping must be relevant to each other. For example, point based spatial grouping with `group_times` may only be relevant with temporal thresholds in units of hours or minutes while line and polygon based spatial grouping with `group_lines` and `group_polys` may only be relevant with temporal thresholds in units of hours or days.

## Generating networks

Here, we will provide an example of point based spatial grouping with spatsoc’s “Newfoundland Bog Cow” example animal telemetry data. The data has a relocation rate of 2 hours but in this case, we consider relocations that occur within 5 minutes of each other. The columns in this data are “id” (character type), “datetime” (character type), “X” and “Y” Universal Transverse Mercator (UTM) coordinates (numeric type). The character type “datetime” will be converted to POSIXct, R’s date time format, the required type for `spatsoc`’s temporal grouping function `group_times`. The coordinates “X” and “Y” must be in a projected coordinate system with units of meters. In this case, the coordinate system is UTM Zone 21 N. We will use a spatial distance threshold of 50 m given the size and behaviour of the study species. The combination of spatial and temporal thresholds means that any individuals within 50 m of each other within 5 minutes will be assigned to the same group.

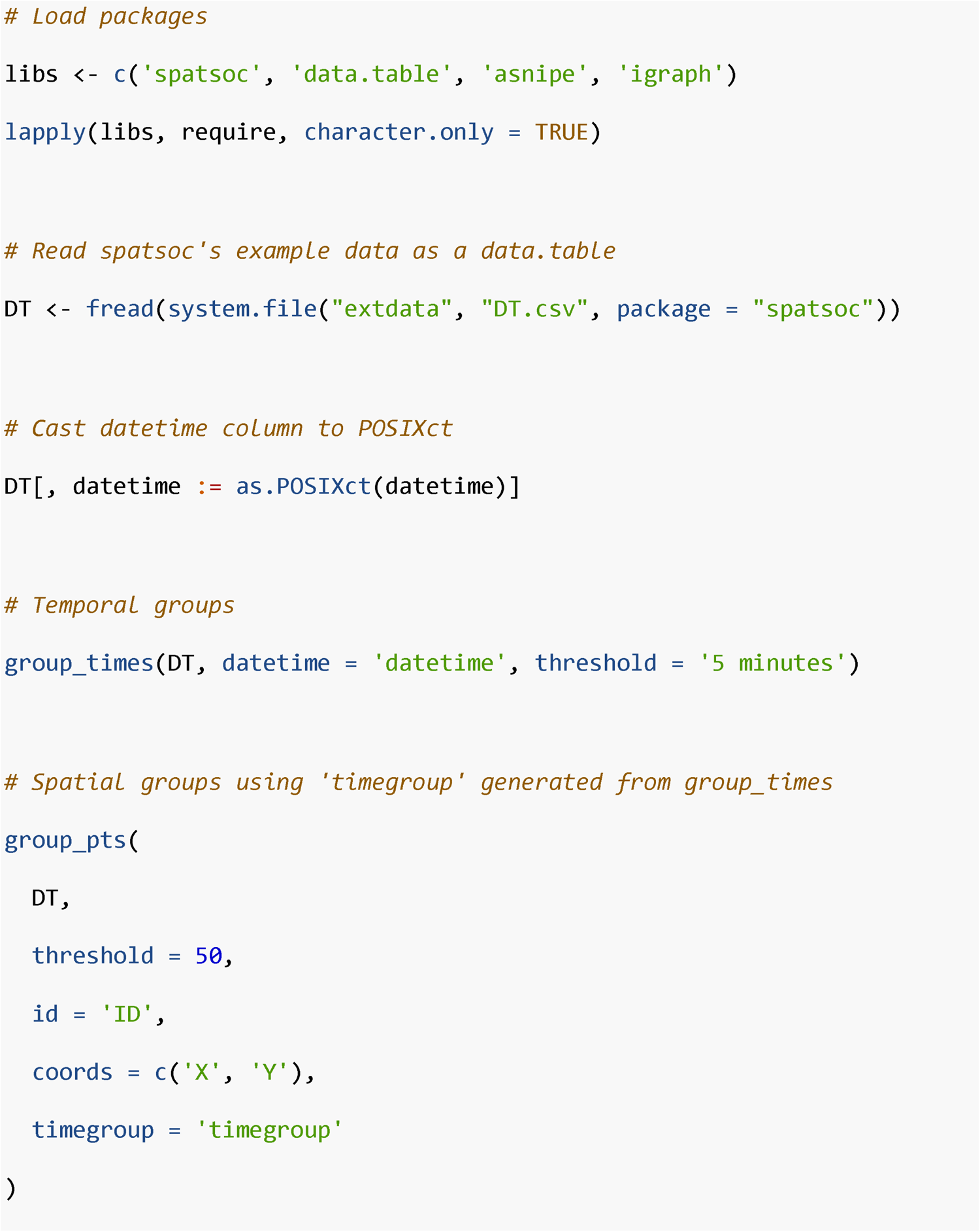

After the temporal and spatial grouping is completed with `group_times` and `group_pts`, a group by individual matrix is generated (described by Farine and Whitehead (2015)). A group by individual matrix forms columns of individuals and rows of groups and a boolean will indicate membership of each individual to a group.

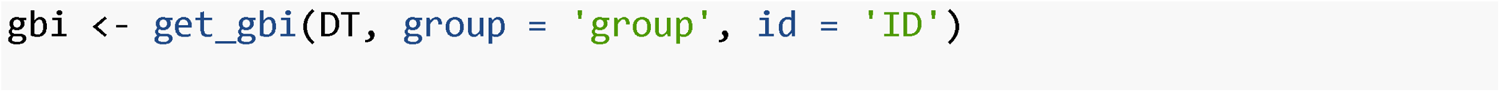

After generating the group by individual matrix, it is passed directly to asnipe, the animal social network package (Farine 2013), to generate a proximity based social network. Note, in this example we use the simple ratio index (SRI) as an association index because all individuals are correctly identified and observed at each relocation event (i.e. the equivalent to an observational period for networks generated using focal observations).

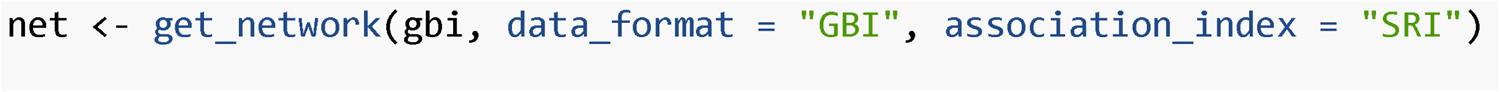

## Data-stream randomization

To perform network data-stream permutations, the `randomizations` function is used to permute spatial and temporal groupings and rebuild PBSNs at each iteration. In this example, we use the “step” method to randomize between individuals at each time step for 500 iterations. The output `randStep` contains the observed and randomized data and can subsequently be used to generate group by individual matrices, networks and calculate network metrics.

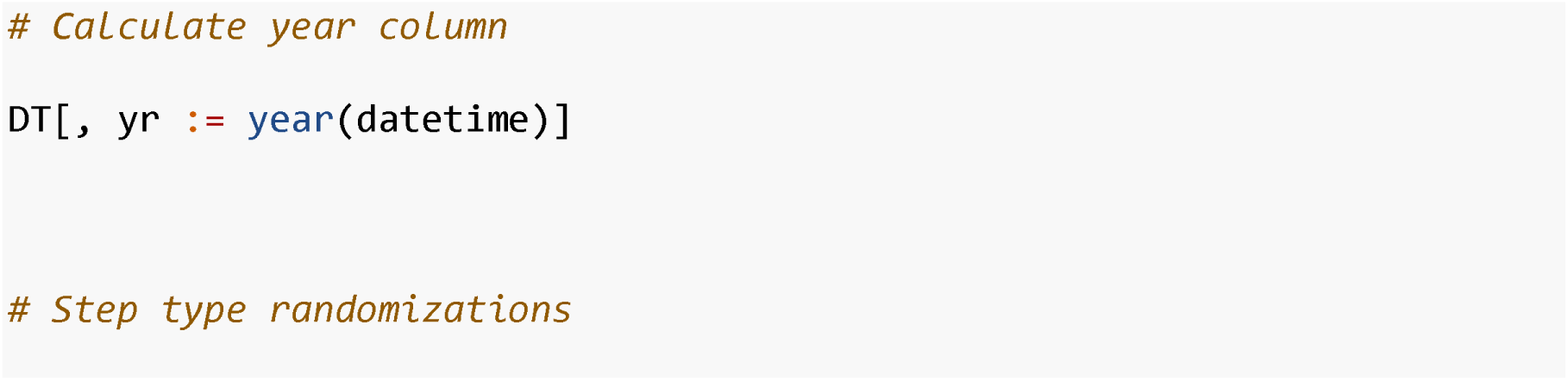
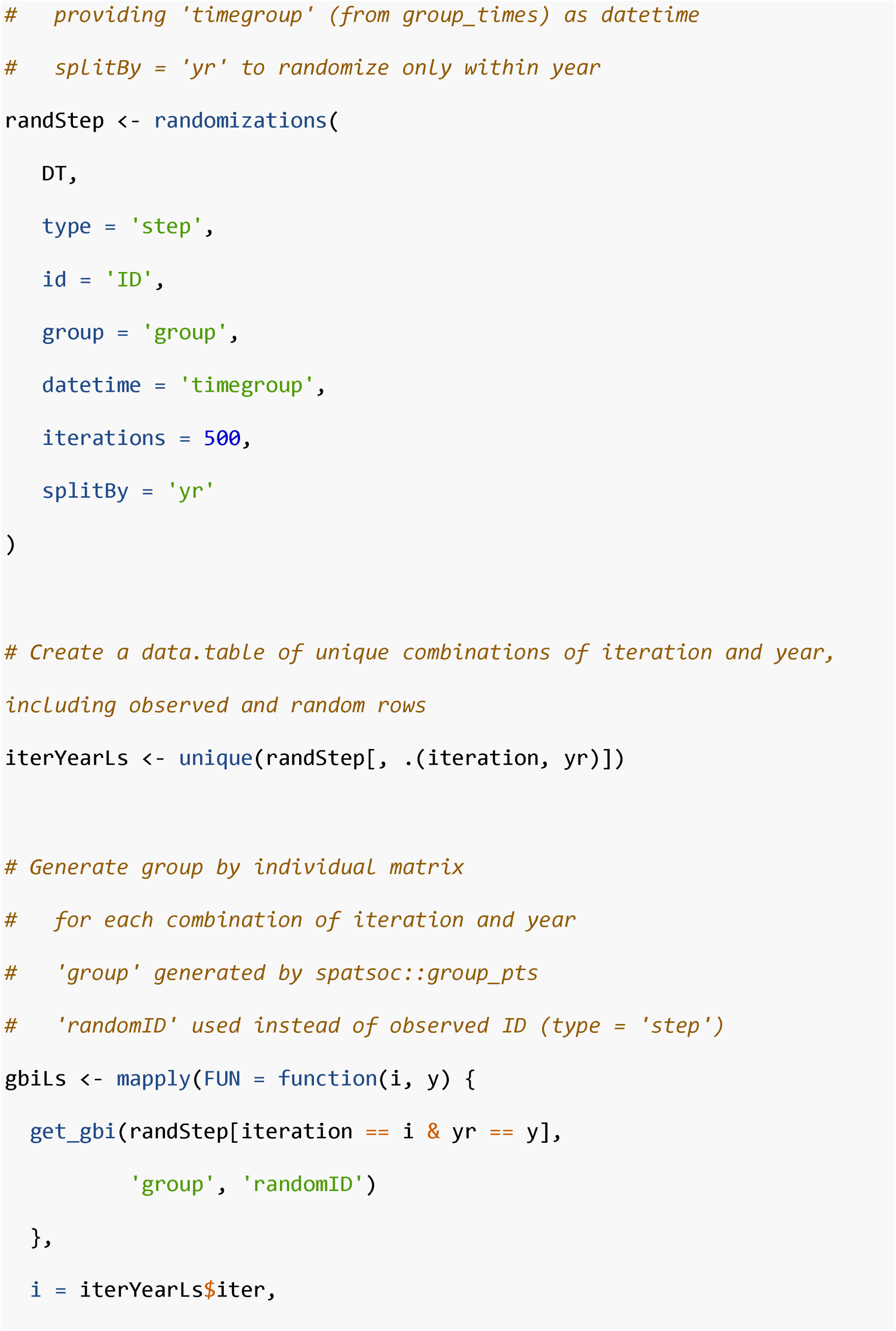
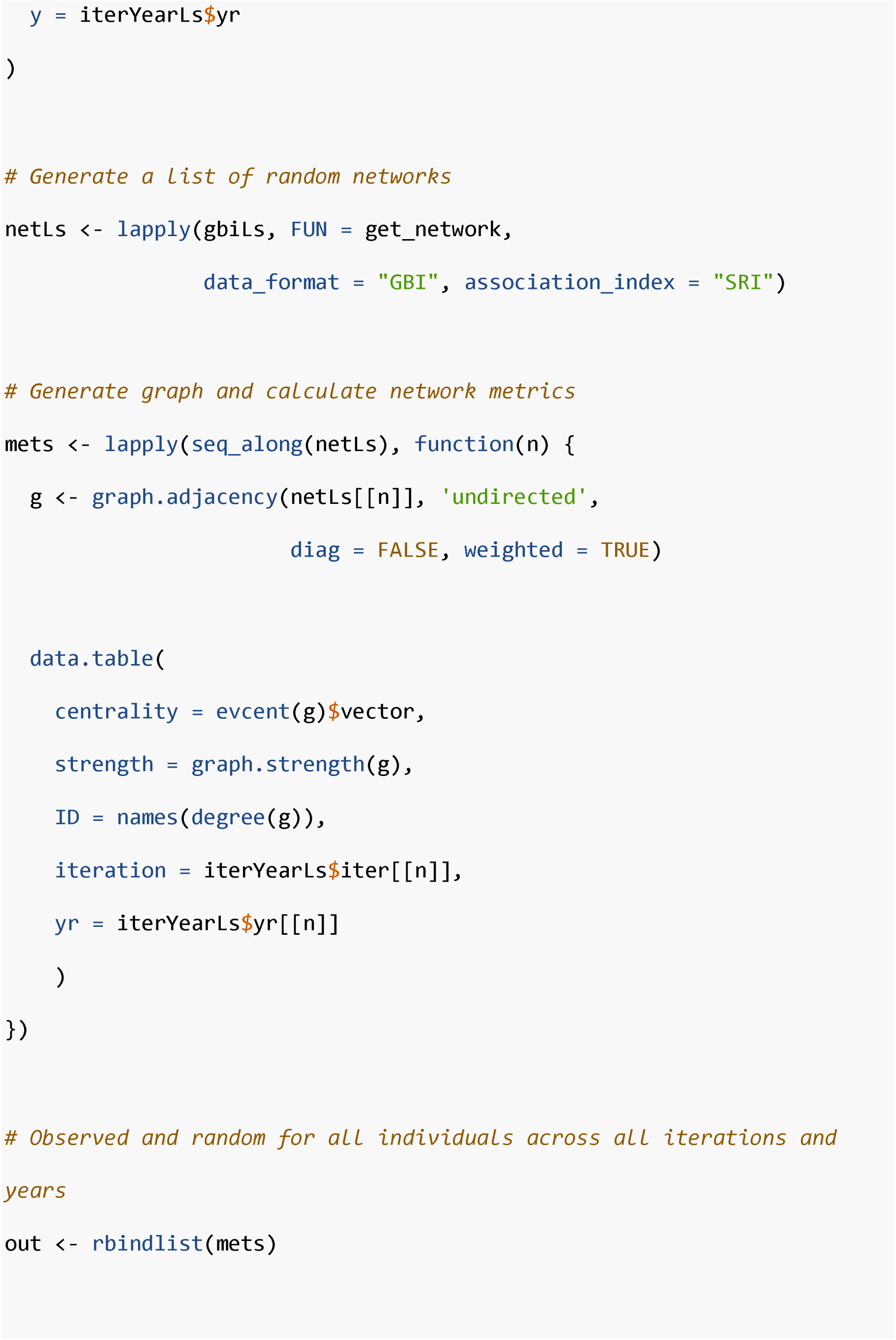
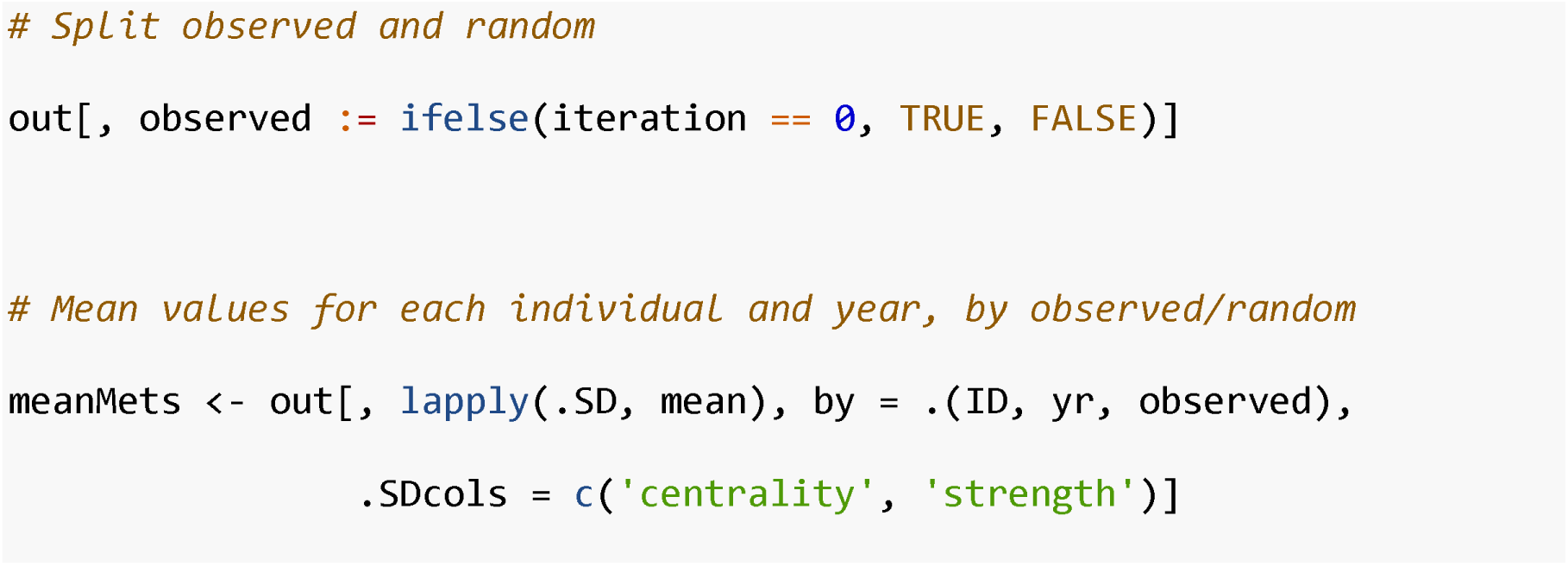

The `splitBy` argument can be used in `randomizations` and grouping functions to specify spatial, e.g. groups or populations, or temporal groups, e.g. weekly, monthly, yearly, by which PBSNs will be generated. For example, in large datasets with individuals in two distinct populations with data over many years, users may use the splitBy argument to generate PBSNs for each population by year combination as opposed to either generating each PBSN separately or using loops.

## Implications

`spatsoc` represents a novel integration of tools for generating PBSNs from animal telemetry data. The grouping and randomization functions allow users to efficiently and rapidly generate a large number PBSNs within the spatsoc environment. `spatsoc` will be of interest and use to a wide range of behavioural ecologists who either already use social network analysis or those who typically work with GPS relocation but are interested in becoming social network users. We advocate for the use of spatsoc in conjunction with other R packages, such as `asnipe` (Farine 2013), `igraph` (Csardi and Nepusz 2006), to facilitate greater sharing of computational and statistical efficiencies and ideas for users of social network analysis.

## Resources

`spatsoc` is a free and open source software available on CRAN (stable release) and at (development version). It is licensed under the GNU General Public License 3.0. `spatsoc` depends on other `R` packages: `data.table` (Dowle and Srinivasan 2018), `igraph` (Csardi and Nepusz 2006), `rgeos` (Bivand and Rundel 2018), `sp` (Bivand, Pebesma, and Gomez-Rubio 2013) and `adehabitatHR` (Calenge 2006). Documentation of all functions and detailed vignettes can be found on the companion website at spatsoc.gitlab.io. Development of `spatsoc` welcomes contribution of feature requests, bug reports and suggested improvements through the issue board at https://gitlab.com/robit.a/spatsoc/issues}.

## Acknowledgements

We thank all members of the Wildlife Evolutionary Ecology Lab, including Juliana Balluffi-Fry, Sana Zabihi-Seissan, Erin Koen, Mike Laforge, Christina Prokopenko, Julie Turner, Levi Newediuk and Chris Hart for their comments on previous versions of this manuscript. We thank Tyler Bonnell, Martin Leclerc and Shane Frank for testing the package ahead of its release. We also thank the rOpenSci organization for their package onboarding process including rOpenSci reviewers for their code review, which contributed to improving this package. Funding for this study was provided by a Vanier Canada Graduate Scholarship to QMRW and a NSERC Discovery Grant to EVW.

## Author contributions

ALR, QMRW, and EVW conceived of the original package concept. ALR developed the package. ALR and QMRW drafted the manuscript and all co-authors contributed critically to the drafts and gave final approval for publication.

## Data accessibility

The data used for illustration is distributed with the package as example data. After installing the package, the data can be viewed with:

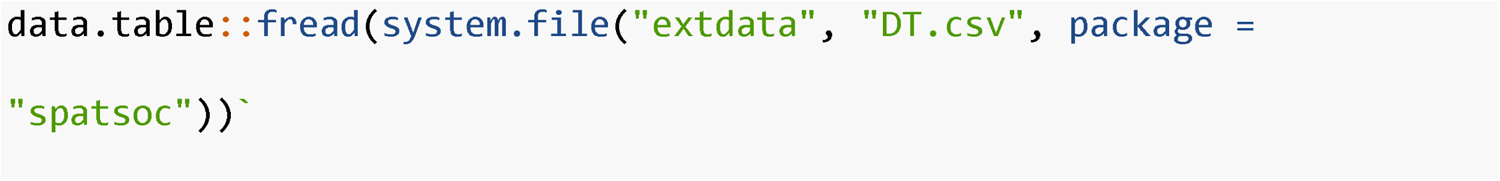

## Citation

Users of `spatsoc` should cite this article directly. A formatted citation and BibTex entry is provided in:

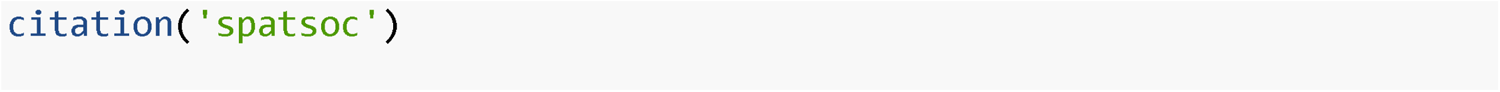

